# Cortical interneurons: fit for function and fit to function? Evidence from development and evolution

**DOI:** 10.1101/2023.02.23.529671

**Authors:** Joram Keijser, Henning Sprekeler

## Abstract

Cortical inhibitory interneurons form a broad spectrum of subtypes. This diversity suggests a division of labour, in which each cell type supports a distinct function. In the present era of optimisation-based algorithms, it is tempting to speculate that these functions were the evolutionary or developmental driving force for the spectrum of interneurons we see in the mature mammalian brain. In this study, we evaluated this hypothesis using the two most common interneuron types, parvalbumin (PV) and somatostatin (SST) expressing cells, as examples. PV and SST interneurons control the activity in the cell bodies and the apical dendrites of excitatory pyramidal cells, respectively, due to a combination of anatomical and synaptic properties. But was this compartment-specific inhibition indeed the function for which PV and SST cells originally evolved? Does the compartmental structure of pyramidal cells shape the diversification of PV and SST interneurons over development? To address these questions, we reviewed and reanalysed publicly available data on the development and evolution of PV and SST interneurons on one hand, and pyramidal cell morphology on the other. These data speak against the idea that the compartment structure of pyramidal cells drove the diversification into PV and SST interneurons. In particular, pyramidal cells mature late, while interneurons are likely committed to a particular fate (PV vs. SST) during early development. Moreover, comparative anatomy and single cell RNA-sequencing data indicate that PV and SST cells, but not the compartment structure of pyramidal cells, existed in the last common ancestor of mammals and reptiles. Specifically, turtle and songbird SST cells also express the *Elfn1* and *Cbln4* genes that are thought to play a role in compartment-specific inhibition in mammals. PV and SST cells therefore evolved and developed the properties that allow them to provide compartment-specific inhibition before there was selective pressure for this function. This suggest that interneuron diversity originally resulted from a different evolutionary driving force and was only later co-opted for the compartment-specific inhibition it seems to serve in mammals today. Future experiments could further test this idea using our computational reconstruction of ancestral Elfn1 protein sequences.

## 1 Introduction

Cortical inhibitory interneurons are a highly diverse group, differing in their morphology, connectivity, and electrophysiology [1]. Decades of experimental and theoretical work have suggested a role for interneurons in many functions [1, 2, 3], including the regulation of neural activity [4, 5], control of synaptic plasticity [6, 7], increasing temporal precision [8, 9], predictive coding [10, 11], and gain modulation [12, 13]. Many of these functions come down to the control of excitation.

Why would the control of excitation require a diversity of interneurons? A key reason could lie in the complexity of excitatory cells [14, 15]. Pyramidal cells (PCs) consist of several cellular compartments that have different physiological properties (e.g. sodium vs. calcium spikes [16]), receive different inputs (e.g., top-down vs. bottom up [17, 18], although see [19]) and follow distinct synaptic plasticity rules [20, 21, 22]. The control of different pyramidal cell compartments might therefore require inhibition from designated types of interneurons. Indeed, the two most common interneuron types—parvalbumin (PV)- and somatostatin (SST)-expressing cells— are classically distinguished by their connectivity with pyramidal cells: whereas PV-expressing basket cells mainly target the somata of PCs, SST-expressing Martinotti cells mainly target their apical dendrites [1]. The cellular and synaptic properties of these interneurons also seem adapted to this purpose. SST interneurons receive facilitating synapses from PCs [23, 24], rendering them sensitive to bursts of action potentials [25, 26, 27] triggered by plateau potentials in the apical dendrite of PCs [16, 28]. Indeed, SST interneurons control dendritic excitability and bursting of PCs [26, 29, 30]. PV interneurons, on the other hand, receive depressing synapses [23, 31], rendering them less sensitive to these signals [32]. The presynaptic dynamics of PV and SST interneurons therefore seem particularly well-matched to the physiology of pyramidal cells, although both types also inhibit non-pyramidal cells and other interneurons (see e.g. [33, 34]). These and similar observations have led to the view that interneuron diversity can be understood from a functional perspective, in which the morphology and synaptic and cellular properties of different interneurons are fit to specific functions (Fig. 1a, [2, 14]). Consistent with this idea that interneurons are adapted to control different pyramidal cell compartments, we recently showed that properties (connectivity and short-term plasticity) of PV and SST interneurons emerge when optimising a network model for compartment-specific inhibition (Fig.1b, [15]).

**Figure 1:**
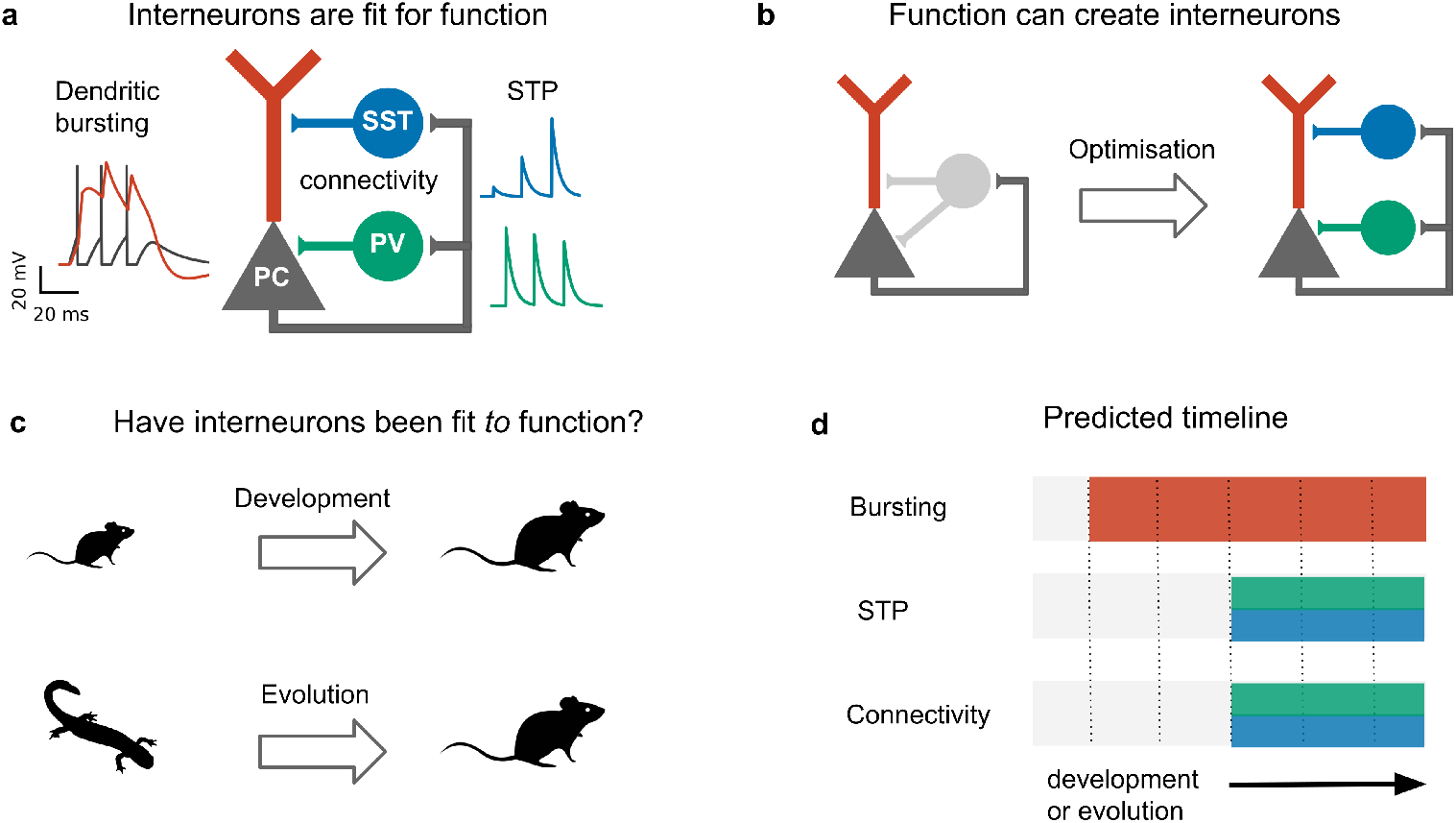
Do development and evolution fit interneurons to function? **(a)** The connectivity and shortterm plasticity (STP) of PV and SST positive interneurons seem adapted to the morphology and electrophysiology of pyramidal cells, as highlighted by an optimization-based model **(b)**. In this model, optimising interneuron parameters to provide compartment-(soma/dendrite) specific inhibition causes interneurons to diversify into two groups that resembled PV and SST interneurons in their connectivity and short-term plasticity [15]. **(c)** The existence of PV and SST subtypes might therefore result from an developmental or evolutionary tuning of interneurons based on pyramidal properties. **(d)** This predicts that immature (or ancestral) circuits contain bursting pyramidal neurons and undiversified interneurons. Development (or evolution) then diversifies the interneurons into PV and SST subtypes. PC activity and short-term plasticity simulated with models from refs. [35, 36] and [37], respectively. Animal silhouettes from https://beta.phylopic.org/

The specialisation of PV and SST interneurons to pyramidal soma and dendrites, respectively, makes it tempting to speculate that the diversification of these interneuron subtypes was driven by pyramidal cell properties, either during evolution or during development (Fig. 1c). This hypothesis predicts a specific temporal order: During evolution or development, the compartmentalization of pyramidal cells should predate interneuron diversification (Fig. 1d).

Here, we evaluate this idea, with a focus on PC and interneuron properties that seem particularly well-adapted to each other: the active dendrites of pyramidal cells, and the connectivity and short-term plasticity of interneurons. Reviewing and re-analysing recent evolutionary and developmental data, we reconstruct the developmental and evolutionary history of these three properties. We find no support for the idea that interneurons develop or evolved to control preexisting compartments of pyramidal cells. Instead, the central properties of PV and SST interneurons that led to this idea emerge before the PC properties they seem adapted to, in both development and evolution. Rather than pyramidal physiology driving interneuron diversification, this suggests a model in which existing interneuron properties enabled new pyramidal cell functions.

## 2 Developmental trajectory of compartment-specific inhibition

We first discuss the developmental trajectory of pyramidal cells and PV and SST interneurons in the mammalian cortex, to assess whether the diversification of PV and SST interneurons during development is driven by pyramidal cell properties. We mostly consider data from rodents, but many of the findings seem to be conserved among mammals [38, 39, 40, 41].

In contrast to pyramidal cells, interneurons are not born in the developing cortex, but subcortically ([42, 43], Box 1). It is only upon migrating to the cortex that interneurons acquire their mature morphology and physiology. The long period between interneuron birth and maturation has led to different models of interneuron development [2, 44]. One model attributes the late maturation of interneurons to a late specification of their cellular identity, possibly based on external cues within the circuit they embed themselves in [2, 44]. Alternatively, the late emergence of characteristic features could be due to the slow unfolding of a predetermined genetic program that happens independently of the surrounding circuit[45].

### Box 1 Birth and migration of cortical interneurons

Cortical GABAergic interneurons are born in a transient region of the developing brain known as the ganglionic eminence [42, 45, 44], from where they tangentially migrate to the cortex [50]. The ganglionic eminence can be divided into multiple subregions patterned by unique combinations of transcription factors [43, 38, 39, 51], that activate distinct genetic programs. Since each genetic program corresponds to a different cell type, the majority of the cells born in the medial ganglionic eminence (MGE) will become PV and SST interneurons, whereas the caudal ganglionic eminence (CGE) generates, among others, vasoactive intestinal peptide (VIP)-expressing interneurons [52, 53, 54, 55, 56]). A key example for a patterning transcription factor that shapes interneuron identity is *Nkx2*-*1*, which is expressed within the MGE but not CGE [57, 55]. *Nkx2*-*1* knockout leads MGE-derived interneurons to adopt the fate of CGE-derived interneurons [58]. Molecular gradients have also been shown to contribute to interneuron diversity within the same eminence: the dorsal-caudal and rostral-ventral MGE preferentially generate SST and PV neurons, respectively [59, 60, 61, 62, 51, 63].

After birth, interneurons migrate to the developing cortex via two different routes: The superficial marginal zone (the MZ, which will develop into cortical layer 1) and the deeper subventricular zone (SVZ). These different migration routes are used by distinct layer 2-3 (L2-3) SST subtypes [45]. Whereas L2-3 SST Martinotti cells migrate via the marginal zone from where they descend to their future location, non-Martinotti cells migrate via the subventricular zone [64]. This indicates that SST subtypes (at least in L2-3) are predetermined before their arrival in cortex, possibly even before they “choose” one migratory route over the other. Future L2-3 Martinotti cells forced to migrate via the wrong route (the SVZ) still become Martinotti cells in terms of their transcriptional profile and electrophysiology, but they lack a fully developed layer 1 axon [64]. This suggests that developing L2/3 Martinotti cells cannot send their developing axon from deeper to upper layers, but have to leave it there while their cell body descends. Translaminar axons of other neurons such as a less studied PV subtype [64], and cerebellar granule cells [65] are established via a similar mechanism, suggesting it might be the only reliable way for neurons to develop translaminar projections.

The malleability of interneuron properties during development is therefore currently an open question: Which properties are adapted to the surrounding circuit, and which are predetermined? Whatever properties are adapted, cellular identity (e.g., PV vs. SST) is probably not one of them [44, 45]. Interneuron types— at least on a coarse level—are determined by their time and place of birth. Future PV and SST interneurons, for example, are preferentially generated within different parts of the same embryonic structure ([46, 45],Box 1).

Recent data suggests that not just interneuron types (e.g., PV vs. SST), but also interneuron *sub*types (e.g., SST Martinotti vs. SST non-Martinotti) are specified early in development. Lim et al. [45] showed that Martinotti and non-Martinotti cells migrate to the developing cortex via different routes (Box 1). In addition, a developing interneuron’s transcriptional profile can be used to predict its future fate [47, 48, 40, 49].

Although interneurons are therefore likely hardwired to become a certain subtype, it is still possible that interneuron properties such as short-term plasticity or connectivity are subject to activity-dependent fine-tuning. For example, the development of short-term facilitation or a layer 1 axon of SST Martinotti cells might emerge in dependence on pyramidal neuron bursting. In this case, bursting should develop ahead of these SST features.

When do developing pyramidal cells first show dendrite-dependent bursting? Their electrophysiology matures relatively late: dendritic plateau potentials emerge only in the third postnatal week [66, 67]. This is consistent with the late maturation of their dendritic morphology. PCs develop their intricate apical arborization and tuft dendrites after the second postnatal week [66, 68]. For example, the tuft length increases almost twofold during the third postnatal week [68], and dendritic spikes fail to reach the soma on postnatal day 14 and 28 [66].

When does short-term facilitation (STF) of PC→SST synapses arise during development? Could its development be driven by bursting in pyramidal cells? Some of the early experiments showed such STF in rat cortex during the third postnatal week [69, 24, 23]. Evidence for an even earlier presence of STF in these synapses comes from molecular studies. The Gosh laboratory has shown that the short-term facilitation in hippocampal [70] and cortical [71] PC→SST synapses is due to the transmembrane protein Elfn1, which is expressed by SST mouse and human interneurons (Box 2, Fig. 2 In these experiments, STF was measured in the second postnatal week, and the expression of Elfn1 was detected already one week after birth [72, 73], providing an early molecular signature of short-term facilitation in SST cells. Short-term facilitation in PC→SST synapses is therefore present before dendrite-dependent bursting in PCs.

**Figure 2:**
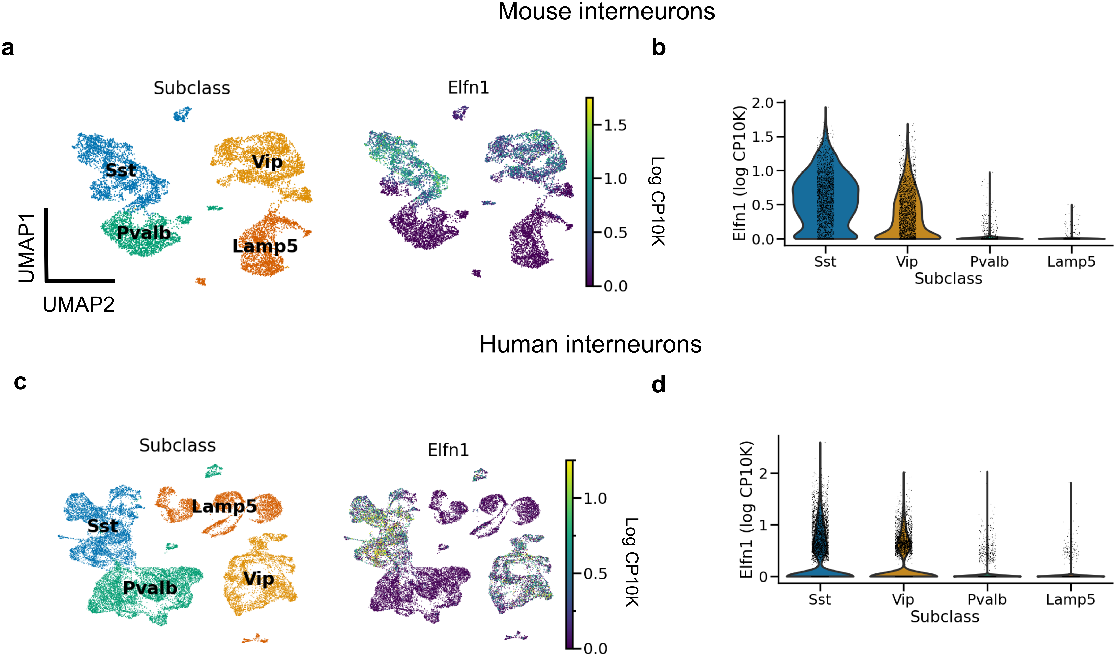
*Elfn1* expression correlates with short-term facilitation in mammals. **(a)** UMAP [77] plot of mouse interneurons coloured by subclass (left) and *Elfn1* expression (right). The two interneuron types — SST and VIP interneurons — known to receive facilitating synapses both express *Elfn1*. **(b)** Violin plot of *Elfn1* expression by subclass. CP10K: counts per 10 thousand. **(c,d)** As **(a,b)**, but for human interneurons. Data from from refs. [74] (a,b), and [75] (c,d).

### Box 2 Genetic basis of short-term facilitation

Pyramidal cells form short-term depressing synapses onto PV neurons, but short-term facilitating synapses onto SST neurons. This difference is partly attributed to the postsynaptic expression of Elfn1 by SST neurons [70, 72, 71]. Elfn1 is a synaptic protein that contacts the presynaptic boutons of pyramidal cells and controls their release properties. Specifically, Elfn1 induces presynaptic localization of metabotropic glutamate receptor 7 (mGluR7) [72]. *Grm7*, the gene coding for mGluR7, is near-ubiquitously expressed in mouse (and human) neurons (data from [74, 75]). mGluR7 has a low affinity for glutamate: Only high glutamate levels caused by repeated presynaptic stimulation will lead mGluR7 to activate calcium channels, which increase synaptic release and thereby mediate synaptic facilitation. Elfn1 causes facilitation of PC→SST synapses in the hippocampus and different cortical layers [71]. As expected from their expression of Elfn1, human SST (and VIP) interneurons receive facilitating inputs ([34], Fig. 2) However, in the mouse brain the correlation between the short-term facilitation and the expression of Elfn1 is very high, but not perfect [76].

What about the second difference between PV and SST neurons, their compartment-specific output synapses? SST and PV cells form compartment-specific synapses in visual cortical organotypic cultures that lack external inputs [78]. This strongly suggests a role for genetic encoding rather than experience-dependent activity. Indeed, recent work identified important molecular players in the establishment of compartment-specific synapses [73]. Several genes are involved in the formation of compartment-specific synapses. For example, suppressing *Cbln4* in SST interneurons decreased inhibition onto PC dendrites. An over-expression of the same gene in PV interneurons, on the other hand, increased inhibition onto PC dendrites [73]. Other genes contribute to somatic inhibition in a seemingly analogous way [73]. Both loss and gain of function were shown around P14. Similarly, somatic inhibition in CA1 abruptly emerges at the end of the second postnatal week [79]. It is therefore by the second postnatal week that PV and SST interneurons are committed as to where to direct their output synapses.

Intriguingly, *Cbln4* is only expressed in a subset of neurons (Fig. 3). Clustering revealed that these *Cbln4* + neurons correspond to previously identified subtypes. The *Tac1* cluster labels non-Martinotti cells that target the dendrites of L4 cells [80, 81, 82], and the *Calb2* and *Etv1* clusters correspond to fanning-out Martinotti cells [80, 5], consistent with a role of *Cbln4* in establishing dendritic synapses. But only a subset of the *Myh8* cluster—corresponding to T-shaped Martinotti cells [5]— expressed *Cbln4*, suggesting diverse mechanisms for dendritic targeting.

**Figure 3:**
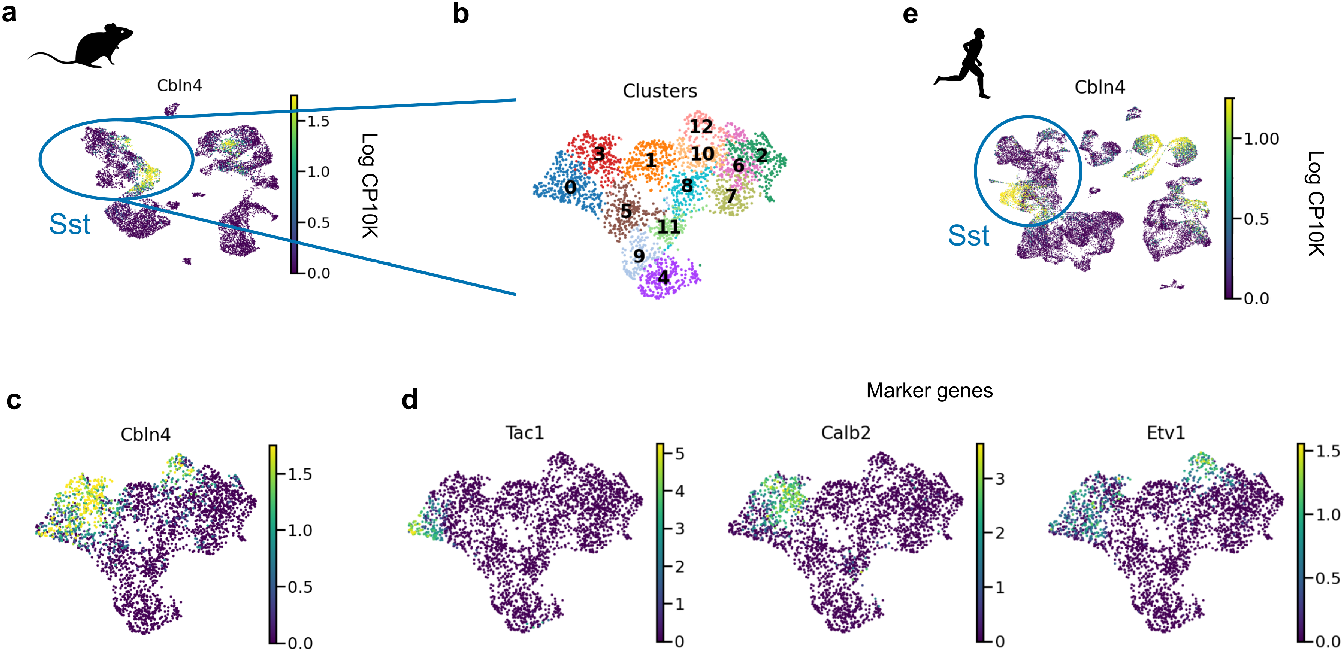
*Cbln4* is expressed in a subset of mammalian SST interneurons. **(a)** UMAP plot of mouse and human interneurons, coloured by their expression of *Cbln4*, a gene that instructs synapse formation onto pyramidal dendrites in mice [73]. *Cbln4* is expressed in certain mouse and human interneuron subtypes, including SST cells. **(b)** UMAP of SST cells, clustered into subgroups. **(c)***Cbln4* is expressed in clusters 0, 3, and 12, which also express marker genes *Tac1*, *Calb2* and *Etv1* **(d)**, respectively. **(e)** A subset of human Sst cells also express *Cbln4.* Data from from refs. [74] (a) and [75] (e).

An interneuron’s cell type, the plasticity of their input synapses from PCs, and the PC compartments they target are therefore determined before interneurons are fully embedded within cortical circuits, and before pyramidal neurons develop their characteristic morphology and electrophysiology. This suggest that while PV and SST interneurons are fit for the function of compartment-specific inhibition of PCs, some of their characteristic properties are probably not developmentally driven by PC activity.

### Evolutionary trajectory of compartment-specific inhibition

On a much longer timescale than development, evolution also changes the properties of cell types. This raises the question whether the differentiation of PV and SST interneurons preceded the evolution of the compartmental complexity of pyramidal neurons.

If natural selection tuned PV and SST neurons to pyramidal cell properties, the brains of mammalian ancestors must have contained pyramidal cells with elaborate dendrites, while interneurons were still undifferentiated (Fig. 1c,d). This hypothesis cannot be tested directly since our mammalian ancestors are no longer alive, and their fossils provide no information regarding cell types. We therefore have to infer the evolutionary history of cell types by comparing data from modern-day species (Fig. 4a, [83, 84]). Although many cell type-specific properties such as short-term plasticity have not been measured in non-standard model organisms (see refs. [85, 86, 34] for recent exceptions), transcriptomic correlates can be studied using single cell RNA sequencing (scRNA-seq, [87]), offering a means for defining and comparing cell types across species [88, 84].

**Figure 4:**
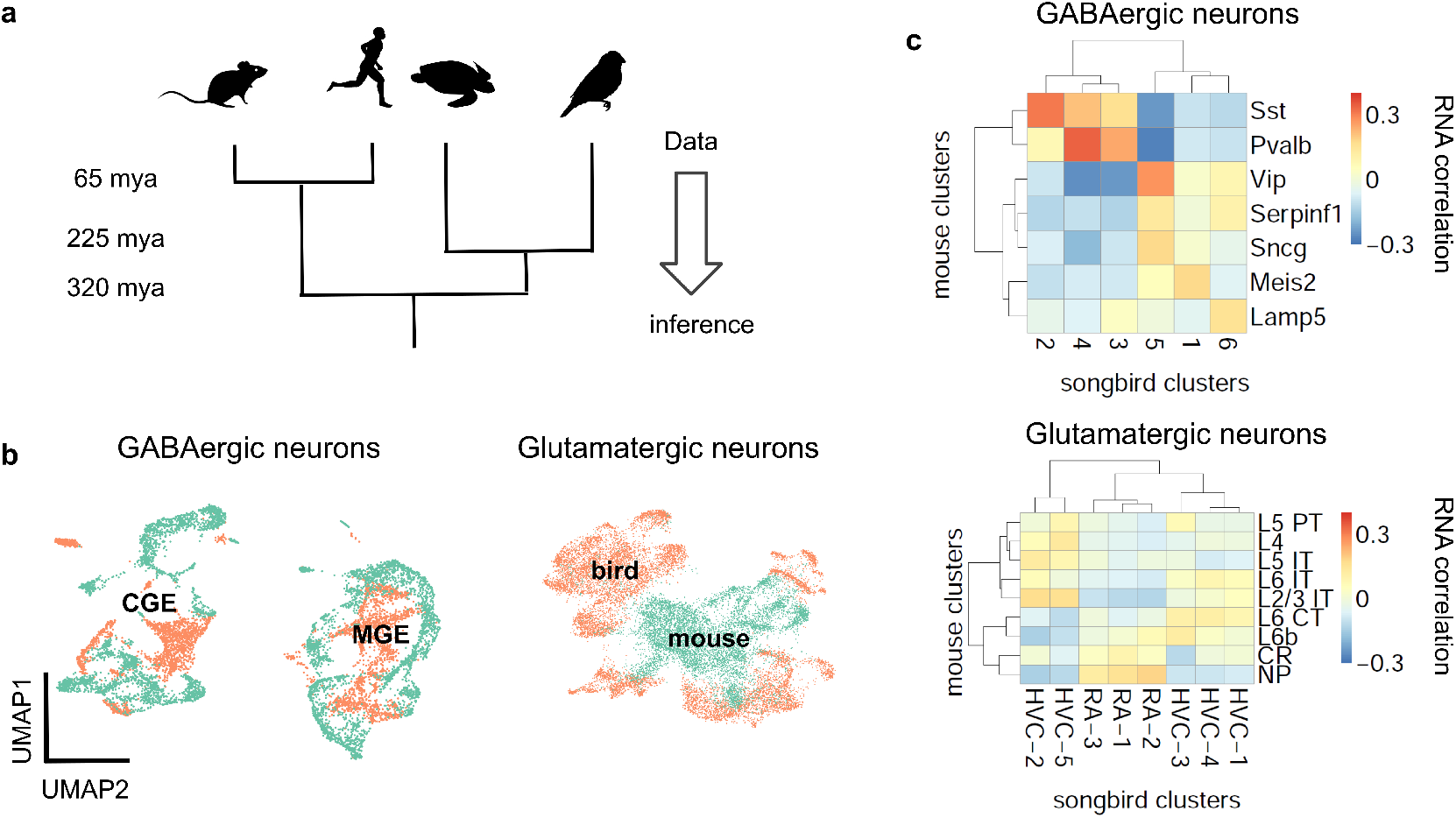
Evolutionary conservation of GABAergic cell types. **(a)** Phylogenetic approach. **(b)** Pearson correlation between average RNA expression in clusters of songbird and mouse interneurons. Correlations between GABAergic neurons are typically larger. **(c)** UMAP plots of integrated gene expression data for GABAergic and glutamatergic neurons. GABAergic neurons first cluster by developmental origin (MGE vs. CGE, see Box 1) and then by species. Mouse data from ref. [74], songbird data and correlation analysis from ref. [92].

#### Interneuron conservation & principal neuron divergence

The first applications of scRNA-seq in neuroscience profiled cell types in mice [89, 90]. More recently, scRNA-seq was used to classify neuron types also in reptiles [91] and songbirds [92]. The evolutionary relationships of reptiles, birds and mammals suggest that a feature found in all three lineages predates their divergence, while a feature found exclusively in the mammalian lineage is, in fact, a mammalian invention. This idea enables inferring the evolutionary history of interneurons and pyramidal cells (Fig. 4a).

Let us first consider the general evolutionary trajectory of excitatory and inhibitory cell types. Tosches et al. [91] used scRNA-seq to analyse cells from the turtle and lizard forebrain and compare them with previously published mammalian data [90]. They found that reptilian inhibitory neurons cluster into groups that roughly correspond to mammalian interneuron types [91]. These results extend earlier findings that found similarities between turtle and mammalian interneurons based on marker genes and morphology [93, 94]. Colquitt et al. [92] recently made analogous observations regarding the similarity of songbird and mouse interneurons (Fig. 4b,c). The most parsimonious explanation of this sharing of interneuron types is that similar types already existed in a common ancestor of the three lineages, rather than convergent evolution in three lineages. This homology is likely due to shared developmental origins: the inhibitory interneurons of birds and reptiles are born within the conserved ganglionic eminences. The fact that interneurons of different lineages are homologous does not mean they are identical. For example, the correlation between mouse PV and SST cells and the best matching songbird clusters is 0.37 and 0.31, respectively (Fig. 4b). This is higher than the correlation between the best-matching glutamatergic types (0.19, see below), but lower than between some of the different cell types within the same species (mouse PV and SST cells: 0.58). Mammalian and non-mammalian interneurons, while homologous, therefore have likely undergone lineage-specific adaptations.

In contrast to inhibitory interneurons, excitatory neurons are probably not homologous between reptiles, songbirds, and mammals (Fig. 4b,c; [91, 92]). Excitatory cell types in different species are defined by different combinations of transcription factors. A clear example is given by the *Fezf2* and *Satb2* genes that specify subcortical [95] and callosal [96] projections, respectively, of mammalian pyramidal cells. Strikingly, these genes are mutually repressive in the mammalian neurons, but co-expressed in reptilian neurons [91, 97]. Comparing excitatory neurons in the songbird and the mammalian brain revealed an analogous pattern: Although excitatory neurons in the songbird forebrain express similar genes as their counterparts in mammalian neocortex, these genes are regulated by different transcription factors [92]. Instead, the transcription factors expressed by songbird glutamatergic neurons are similar to those in e.g. the mouse olfactory bulb and olfactory cortex. Since transcription factors specify cellular identity [98, 83] this suggests that excitatory neurons are not conserved across mammals, birds and reptiles [91, 92, 99].

Inhibitory cell types therefore seem more conserved than excitatory cell types, which appears broadly inconsistent with an evolutionary adaptation of interneurons to pyramidal cells. This is further confirmed when considering the evolutionary history of specific features of excitatory and inhibitory interneurons, in particular, elaborate dendrites and dendrite-dependent bursting and short-term plasticity.

#### Evolution of cell type-specific features

We are not aware of direct measurements of short-term facilitation in non-mammalian species and therefore aimed to infer its presence from the expression of *Elfn1* (Box 2). To this end, we re-analyzed publicly available gene expression data for reptilian and songbird interneuron types [91, 92]. We found that *Elfn1* is also expressed in the types corresponding to mammalian SST (and VIP) interneurons (Fig. 5), but not in the type corresponding to PV interneurons. This suggests that SST-like interneurons expressing *Elfn1* — and potentially faciliating glutamatergic input synapses — were already present in the last common ancestor of reptiles, songbirds and mammals. In terms of potential transcriptomic correlates of synaptic specificity, we find that *Cbln4* (Fig. 3) is expressed in certain subtypes of turtle SST neurons (Fig. 6a). Songbird SST neurons, on the other hand, do not express *Cbln4* (Fig. 6b). The expression of *Cbln4* by Sst interneurons therefore correlates with the presence of apical dendrites in pyramidal cells (Fig. 7, see next). The most parsimonious explanation is that *Cbln4* expression was lost in the songbird lineage. Alternatively, it could have evolved independently in the mammals and reptiles.

**Figure 5:**
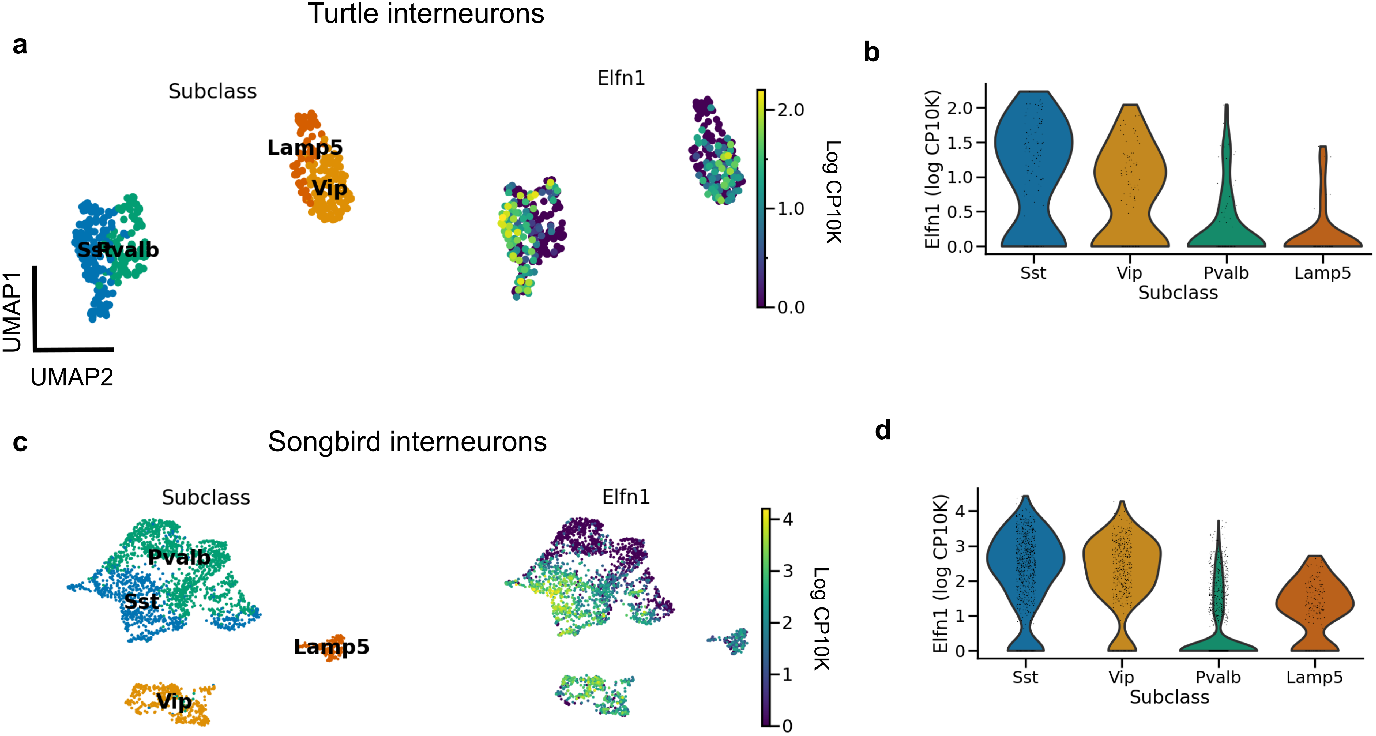
Evolutionary conservation of *Elfn1* expression. **(a)** UMAP plot showing over-expression of *Elfn1* in SST-like and VIP-like interneurons in the turtle forebrain. Data from ref. [91].**(b)** Violin plots of *Elfn1* expression for each of the clusters. **(c,d)** As (a,b), but for zebra finch neurons. Data from ref. [92].

**Figure 6:**
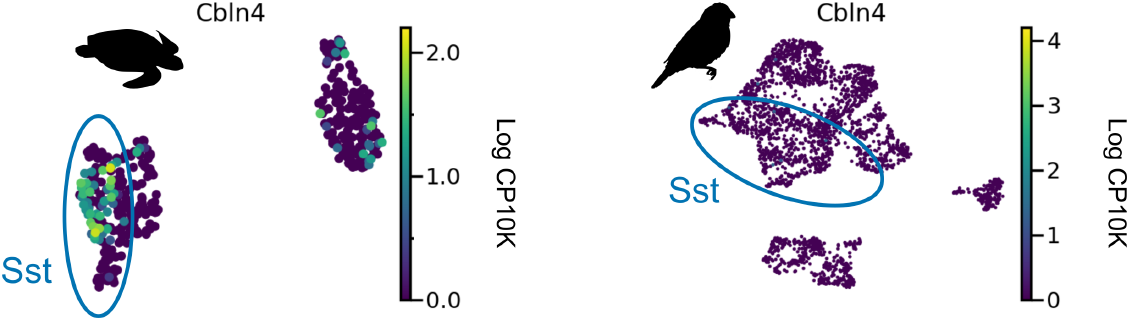
*Cbln4* expression in non-mammalian species. *Cbln4* is expressed in certain subtypes of turtle SST neurons, but not in songbird SST neurons. Data from refs. [91, 92].

**Figure 7:**
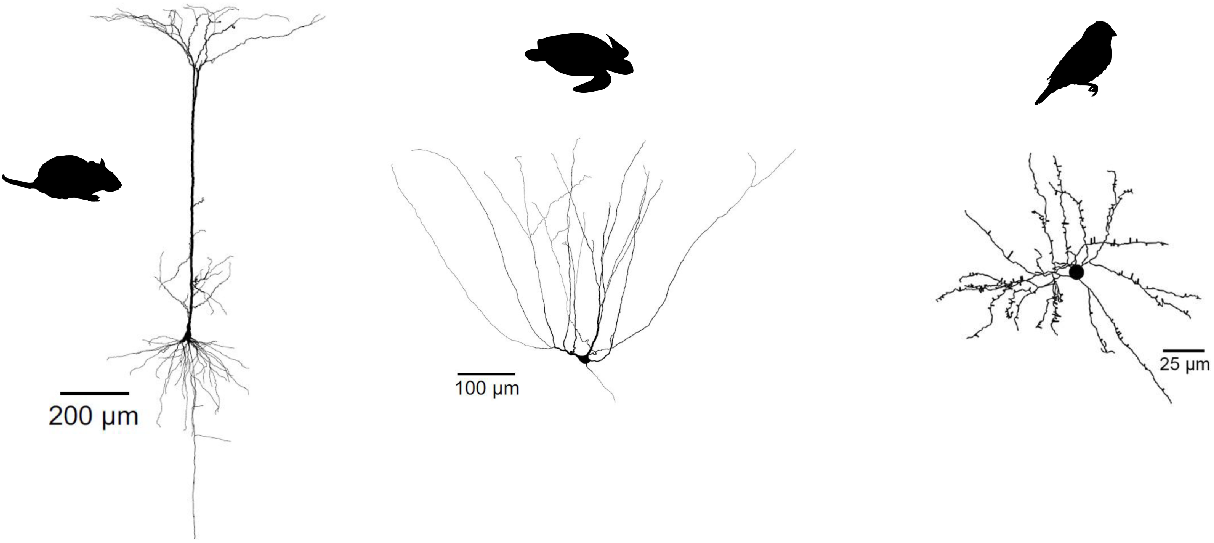
Evolutionary divergence of projection neuron morphology. Both turtle and mammalian projection neurons have a pyramidal morphology, but only mammalian pyramidal neurons have a single apical dendrite. Songbird projection neurons have a stellate, not pyramidal morphology. Turtle and mammalian neurons adapted from [101] (published under a Creative Commons License [107]). Songbird neuron adapted from [108] (published under a Creative Commons License [109])

So not just interneuron subtypes, but also some of their specific properties seem evolutionarily conserved. In contrast, glutamatergic cell types in reptiles and birds show a very different dendritic morphology and physiology from their mammalian pyramidal counterparts. Turtle pyramidal cells have multiple apical dendrites, but no basal dendrites (Fig. 7, [100]). This clear morphological difference suggests that turtle pyramidal neurons are also electrophysiogically distinct. Larkum et al. [101] showed that, *in vitro*, turtle pyramidal neurons lack dendritic calcium spikes and dendrite-dependent bursting. Morphologically similar pyramidal cells in rodent piriform cortex also lack active dendrites ([102, 103], but see [104]). Interestingly, this is probably not due to an absence of calcium channels, but rather to the presence of A-type potassium channels [103]. Songbird excitatory cells have a stellate morphology, and differ therefore even more from mammalian pyramidal cells (Fig. 7, see e.g. [105, 106]).

The lack of dendrite-dependent bursting in reptiles and songbirds is consistent with comparative electro-physiology within the mammalian brain. Pyramidal neurons in the piriform cortex are homologous to certain types of glutamatergic turtle and songbird neurons [92], and also lack dendritic plateau potentials [102]. Pyramidal neurons with mammalian electrophysiological properties therefore evolved after interneurons differentiated into PV and SST cell types. (Fig. 8).

**Figure 8:**
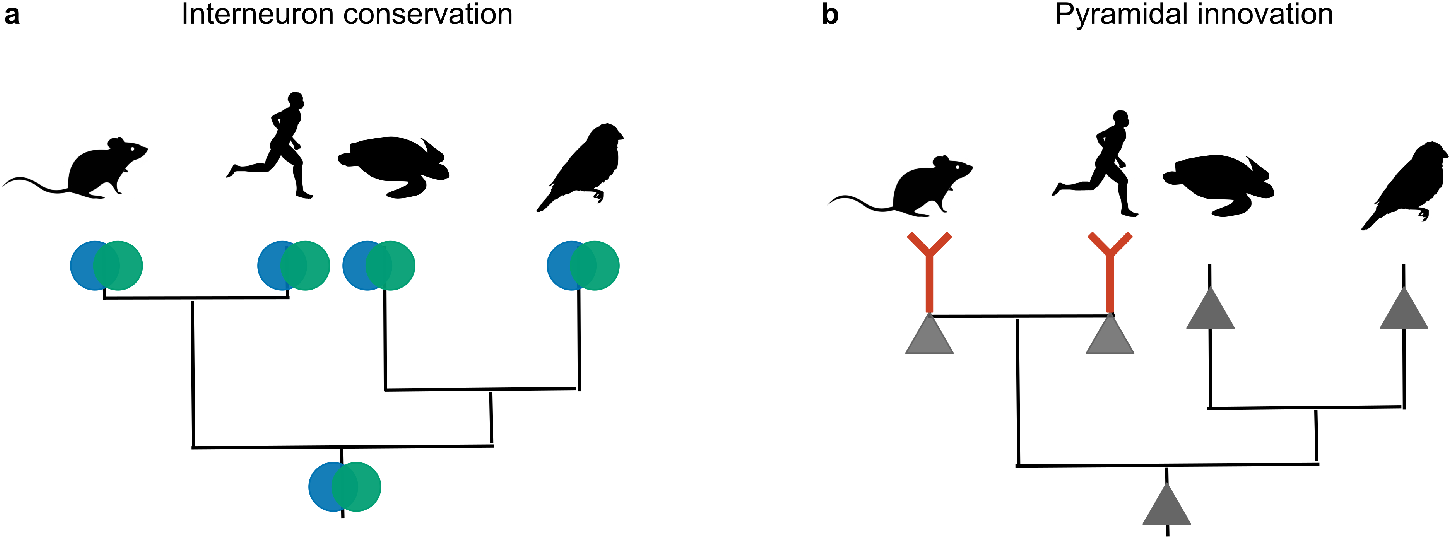
Phylogenetic inference of interneuron and pyramidal evolution. **(a)** mice, humans, songbirds and turtles all have PV and SST interneurons. The most likely explanation for these similarities is that the interneuron types were already present in the last common ancestor of these lineages. **(b)** Only mammalian glutamateric neurons are known to exhibit dendritic plateau potentials that can elicit burst firing. Other lineages probably lack this trait. The most likely explanation is that dendritic bursting evolved only once, in the mammalian lineage.

#### Ancestral Elfn1 reconstruction

The expression of *Elfn1* by zebra finch and turtle SST-like neurons suggests these cells—and therefore the ancestral SST-like cells— receive(d) facilitating inputs. But it is also possible that the ancestral Elfn1 protein had different functional properties. Previous work has used Elfn1 knockout [70, 110, 72] and targeted deletions [111] to discover functionally important domains of the mouse variant (Fig 9b). To determine the evolution of Elfn1 at the resolution of individual sites, we computationally reconstructed its ancestral state ([112, 113], see Methods), starting from the protein sequences of extant species (Fig. 9a). Alignment of the extant sequences revealed that on average across species 74.6% of the Elfn1 sites was identical to that of the mouse protein (Fig. 9c). Combining the sequence alignment with a probabilistic model of sequence evolution [114] and a phylogenetic species-tree allowed us to reconstruct the ancestral protein (Fig. 9d). The amount of conservation varied between protein domains and extant species: the zebra finch and turtle sequences were more similar to the ancestral sequence than the mammalian sequences (Fig. 9e). Future work could use the reconstructed sequences to determine the evolutionary history and the molecular mechanisms of short-term facilitation.

**Figure 9:**
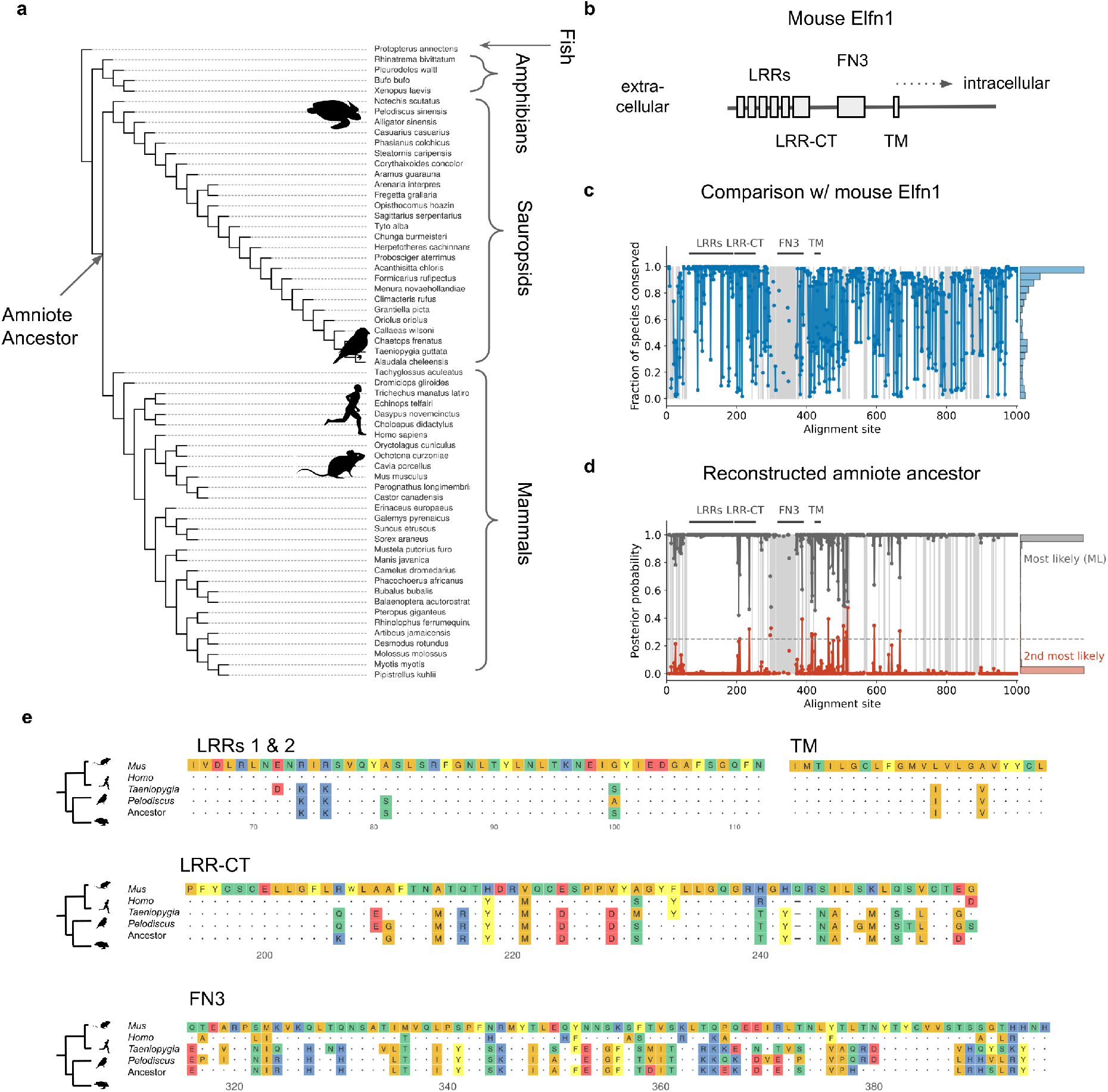
Reconstruction of ancestral Elfn1 protein. **(a)** Species-tree showing the phylogenetic relationships of the species whose Elfn1 homologues were used to reconstruct the Elfn1 protein of the amniote ancestor. **(b)** Domain structure of mouse Elfn1 [115, 111]. LRR: leucine-rich repeat; CT: C-terminal domain; FN3: fibronectin type 3 domain; TM: transmembrane domain. **(c)** Per-site conservation across the tree shown in (a), computed as the fraction of extant species that share the mouse amino acid in a given site. Dashed lines correspond to gaps. Mean conservation: 0.746. **(d)** Posterior probability of ancestral protein. Gray: most likely (ML) sequence, red: 2nd most likely. Dashed line: cutoff for using the 2nd most likely base in “allAlt” sequence. Mean posterior: 0.986. **(e)** Multiple sequence alignment of protein domains shown in (a). Only the first 2 LRRs are shown for space reasons. Dots indicate identity to mouse site, dashes indicate gaps.

## Discussion

The suspicious match between the synaptic properties of PV and SST interneurons and the postsynaptic pyramidal cell compartments suggests that these interneuron properties could be the result of an adaptation to pyramidal cells. Here, we evaluated this idea of interneurons being ‘fit to function’ from an evolutionary and developmental perspective, and showed that the relevant interneuron properties predate those of pyramidal cells both during development and in evolutionary history.

Two lines of evidence indicate that the development of PV and SST interneurons is not induced by mature pyramidal cell activity. First, interneurons become committed to a particular cell type (e.g., PV or SST) before reaching the developing cortex. Interneuron fate therefore cannot be influenced by the activity of pyramidal cells. Second, at least some of the properties of PV and SST interneurons that strongly shape their control of pyramidal cells — short-term plasticity and output connectivity — emerge before the maturation of pyramidal cell morphology and dendritic activity (dendrite-dependent bursting). It should be noted that other interneuron properties clearly are influenced by pyramidal cell activity. Excitatory activity regulates both the survival of interneurons [116], and the formation of inhibitory synapses [117, 118]. Specific types of excitatory neurons determine the laminar allocation of interneurons [119, 120], and their activity can even change the intrinsic properties of mature interneurons [121]. Cell-extrinsic cues therefore play a role in the normal development of interneurons, but are unlikely to determine their identity and the properties we focused on here.

Analogous arguments suggest that the evolution of PV and SST interneurons also cannot be driven by the dendritic physiology of pyramidal cells. The lineages of birds, reptiles and mammals diverged over 300 million years ago, yet they all contain roughly similar interneuron types — evidence that these types were already present in a common ancestor of the three lineages. In contrast to interneurons, excitatory neurons are not conserved, and therefore probably evolved later. The second line of evolutionary evidence relates to two specific aspects of interneuron diversity: short-term plasticity and output connectivity. Recent scRNA-seq data [91, 92] show that reptilian and songbird SST interneurons express *Elfn1*, the gene that in mouse SST neurons is necessary and sufficient for short-term facilitation. Certain reptilian, but not songbird, SST subtypes also express *Cbln4* that plays a role in the synaptic specificity of mammalian SST cells [73].

These data suggest that ancestral interneurons already comprised PV- and SST-like cell types characterised by some of the genes for cell type-specific phenotypes in mammalian interneurons. It does not, however, imply that these phenotypes were actually present in ancestral cells. The expression of *Elfn1*, for example, is not sufficient for facilitating inputs, as shown in the case of VIP subtypes: Multipolar and bipolar VIP neurons both express *Elfn1*, but only the multipolar subtype receives facilitating excitation [71]. It will therefore be interesting to directly test the presence of PV- and SST-specific phenotypes in reptiles and birds. If neither the reptile nor the songbird homologue of SST interneurons receives facilitating excitatory inputs, *Elfn1* was likely reused for short-term facilitation in mammals. The emergence of short-term facilitation in SST neurons would then be an adaptation to pyramidal bursting, co-opting pre-existing interneuron diversity for “pyramidal cell purposes”. The anatomical connectivity of interneurons might similarly have been reused to control pyramidal cells. In the mammalian brain, PV and SST interneurons inhibit not just the somata and dendrites, respectively, of pyramidal cells but also of non-pyramidal cells. Ancestral PV and SST interneurons might therefore have specialised in compartment-specific inhibition, but not of pyramidal cells for which their presynaptic dynamics are so well-matched.

Although our results show that pyramidal cell bursting is unlikely the driver of the differentiation of PV and SST interneurons, this is not in conflict with the functional interpretation of these cell types. In fact, an evolution of active pyramidal cell dendrites before the presence of specialised interneurons would have resulted in aberrant excitation, as seen, e.g., in *Elfn1* mutants [110, 72]. This suggests an alternative picture, in which excitatory neurons can only evolve in a way that still allows the existing interneurons to regulate their activity. This still leaves open the question why interneuron diversity evolved in the first place, if it was not for compartment-specific inhibition. Although it is possible that the initial separation between PV and SST cell types was selectively neutral, this is unlikely given their evolutionary conservation. Instead, the existence of PV and SST cells presumably offers advantages to mammalian and non-mammalian brains alike. An important example of an conserved function could be the temporal coordination of inputs and outputs of pyramidal cells based on oscillations [122, 123].

Our findings have potential implications for the neuroscientific interpretation of optimisation-based models of neural networks, which have recently seen a renaissance [124, 125, 126, 127, 128]. Most of these models describe neural data at the relatively abstract level of dynamics and representations [129, 130]. Recently, such deep network models have also started to include circuit-level structure such as separate excitatory and inhibitory populations [131, 132], different neuronal timescales [133, 134], and short-term plasticity [135, 15]. Deep learning is therefore gradually making its way down from the level of dynamical systems to that of circuits, potentially revealing functional roles for circuit elements. Our findings highlight a challenge to achieving this goal: Multiple circuit-level features — such as the properties of interneuron and pyramidal cells — are interdependent. The function of one feature might depend on that of another and vice versa, raising the question which features should be optimized (e.g., interneurons), and which should be assumed as pre-exising constraints or opportunities (e.g., nonlinear PC dendrites). In other words, optimization-based models face the challenge of modelling processes such as co-evolution. Merging the functional and evolutionary/developmental perspectives will therefore be an important challenge for future work.

## Acknowledgements

We thank Simon J.B. Butt, Loreen Hertäg and members of the Sprekeler Lab for comments on the manuscript.

## 3 Methods

Code was written in Python (version (v) 3.10.8 [136]) and R (v4.2.1 [137]), based on practices outlined in the Good Research Codebook [138]. Code for the transcriptomic analyses can be found at https://github.com/JoramKeijser/interneuron_evolution [139]. Code for the protein reconstruction can be found at https://github.com/JoramKeijser/elfn1_reconstruction [140]

### Datasets

We analysed the following publicly available single cell RNA sequencing data sets: Mouse data from ref. [74] (downloaded from [141]), human data from ref. [75] (downloaded from [142]), zebra finch data (downloaded from [143]), and turtle data from ref. [91] (downloaded from [144]). The paper’s code repository contains a script for automatically downloading the corresponding files.

For each data set, the starting point of our analysis was a matrix of gene counts per cell, together with the clustering of cells from the original publications. We converted each of the datasets to Seurat (v4 [145]) and AnnData (v0.8 [146]) objects for downstream analysis in Python and R, respectively. For visual comparison, we labeled songbird and turtle cell clusters according to the most similar mammalian interneuron subclass, as determined in the original publications. This involved the merging of fine-level clusters that presumably capture within-subclass differences. For each dataset, we only visualized cells part of, or corresponding to, cortical interneurons. In particular, we did not visualize the correlation of the songbird GABAergic clusters 7, 8, and Pre, since these seem homologous to mouse olfactory bulb interneurons [92].

### Dimensionality reduction and clustering

We used AnnData and Scanpy (v1.9.1 [147]) to visualize the expression of the *Elfn1* and *Cbln4* genes. This was done separately for each dataset. We first scaled the counts from each cell to counts per 10 thousand (CP10K) to account for differences in sequencing depth. We then used log plus one pseudo count (log1p) as variance-stabilizing transformation. Finally, we reduced the dimensionality of each dataset, by first finding highly variable genes, performed PCA followed by UMAP [77]. We used scanpy’s default parameters for each of these steps. To investigate *Cbln4* expression within the SST population, we performed dimensionality reduction on all SST cells except long-range projecting Chodl cells. Clustering was done using the Leiden algorithm [148] with resolution 1.

### Correlation analysis

We quantified the overall similarity of species-specific cell clusters by replicating the correlation analysis from refs. [91, 92]. We separately performed the following analysis on GABAergic and glutamatergic cells, and only compared zebra finch and mice. Specifically, we performed the following steps.

1. Select genes to compare across species. For each species, determine subclass-specific marker genes using Seurat’s findAllMarkers (t-test, min.pct = 0.2, max.cells.per.ident = 200) and retain genes with Bonferroni adjusted p-value below 0.05.
2. Intersect the two species-specific lists to find genes that are differentially expressed in both species. This resulted in approximately 500 genes, depending on the cell type.
3. Average counts within each cluster and transform to log scale for variance-stabilization. Specifically, compute: log(1 + *x*) + 0.1, with *x* the average count.
4. Divide each gene’s value by its average across clusters to obtain a “specificity score” invariant to a genes’ overall expression [91].
5. Compute the Pearson correlation between all pairs of mouse and songbird clusters.

We visualised the result using the R package pheatmap (v1.10.12 [149]).

### Dataset integration

We used Seurat’s anchor-based integration [150] to integrate the zebra finch and mouse data. We did this for GABAergic and glutamatergic neurons separately. First, we jointly performed normalization and variance stabilization for each dataset using Seurat’s scTransform [151], with the percentage of mitochondrial counts as covariate. Next, we found the top 3000 most variable features across datasets, and used these to identify a set of anchors. These were then used to integrate the datasets. Finally, we jointly analysed the integrated datasets using Seurat’s standard visualization pipeline: scaling and centering, PCA, and UMAP.

### Ancestral Elfn1 reconstruction

We used the Topiary pipeline [113] to reconstruct the amino acid sequences of the ancestral Elfn1 protein based on sequences of extant species. To this end, we first constructed a source dataset consisting of the Elfn1 sequences from *Mus musculus* (mouse), *Homo sapiens* (human), *Taeniopygia guttata* (zebra finch), and *Pelodiscus sinensis* (Chinese softshell turtle). Next, we used Topiary’s seed-to-alignment to find sequence homologs, perform reciprocal BLAST [152] to predict their orthology, reduce sequence redundancy, and align the remaining sequences using Muscle5 [153]. This resulted in 62 aligned sequences that were used as input to Topiary’s alignment-to-ancestors. This infers the maximum likelihood (ML) gene tree, the ML substitution model, and the ML ancestral sequences using RAxML-NG [154]. The posterior probability of an ancestral amino acid was computed using the amino acid’s likelihood weighted by its prior probability, normalised by the sum over all amino acids. Topiary generates bootstrap replicates of the ML gene tree, and uses GeneRax [155] to reconcile the gene tree with the species tree. The number of bootstrap replicates was 700, as automatically determined by the software. This number—but not the reconstructed ML sequence—varied slightly between runs. Finally, we used topiary’s bootstrap-reconcile that estimates the branch support for the reconciled tree. The ancestral sequence contained 16 ambiguous sites (based on a posterior probability cutoff of 0.25). Besides the ML sequence, we also report a worst case “altAll” sequence in which these ambiguous sites have been replaced by the next most-likely amino acid. Branch support for the amniote ancestor was 100/100, indicating very high confidence in the existence of this ancestor, as expected. We aligned extant and ancestral sequences using Muscle5, and visualised the resulting alignment using the R package Ggmsa [156].

## Notes

### Competing Interest Statement

The authors have declared no competing interest.

### Summary of Updates

Cosmetic changes to figures

